# Naturally-acquired and Vaccine-induced Human Monoclonal Antibodies to *Plasmodium vivax* Duffy Binding Protein Inhibit Invasion of *Plasmodium knowlesi* (PvDBPOR) Transgenic Parasites

**DOI:** 10.1101/2023.03.07.531647

**Authors:** Quentin D. Watson, Lenore L. Carias, Alyssa Malachin, Karli R. Redinger, Jürgen Bosch, Martino Bardelli, Robert W. Moon, Simon J. Draper, Peter A. Zimmerman, Christopher L. King

## Abstract

The Duffy antigen receptor for chemokines (DARC) expressed on erythrocytes is central to *Plasmodium vivax* (Pv) invasion of reticulocytes. Pv expresses a Duffy binding protein (PvDBP) on merozoites, a DARC ligand, and their protein-protein interaction is central to vivax blood stage malaria. Here we compared the functional activity of humAbs derived from naturally exposed and vaccinated individuals for the first time using easily cultured *P. knowlesi* (Pk) that had been genetically modified to replace its endogenous PkDBP orthologue with PvDBP to create a transgenic parasite, PkPvDBPOR. This transgenic parasite requires DARC to invade human erythrocytes but is not reticulocyte restricted. Using this model, we evaluated the invasion inhibition potential of 12 humAbs (9 naturally acquired; 3 vaccine-induced) targeting PvDBP individually and in combinations using growth inhibition assays (GIAs). The PvDBP-specific humAbs demonstrated 70-100% inhibition of PkPvDBPOR invasion with the IC_50_ values ranging from 51 to 338 μg/mL for the 9 naturally acquired (NA) humAbs and 33 to 99 μg/ml for the 3 vaccine-induced (VI) humAbs. To evaluate antagonistic, additive, or synergistic effects, six pairwise combinations were performed using select humAbs. Of these combinations tested, one NA/NA (099100/094083) combination demonstrated relatively strong additive inhibition between 10-100 μg/mL; all combinations of NA and VI humAbs showed additive inhibition at concentrations below 25 μg/mL and antagonism at higher concentrations. None of the humAb combinations showed synergy. This PkPvDBPOR model system enables efficient assessment of NA and VI humAbs individually and in combination.

**IMPORTANCE:** Given the importance of Duffy blood group antigen and *P. vivax* Duffy binding protein (PvDBP) interaction leading to blood stage vivax malaria, development of vaccines or therapeutic human monoclonal antibodies (humAbs) targeting PvDBP are key strategies for treating and controlling Pv. The *P. knowlesi*-based PkPvDBPOR transgenic model system enables efficient assessment of NA and VI humAbs individually and in combination. As such, this model could prioritize specific humAb combinations ahead of clinical trials of these reagents.

## INTRODUCTION

The estimated annual global burden of *Plasmodium vivax* (Pv) malaria is 14.3 million (13.7 to 15.0 million) cases (1). However, this approximation of Pv clinical cases grossly underestimates Pv asymptomatic or latent infections in the liver, leading to more subtle morbidity and death in impoverished settings where endemic populations frequently experience malnutrition, co-infections, and limited access to health care (1–3). Pv infections can also include Duffy-negative individuals in sub-Saharan Africa, previously considered protected from Pv erythrocytic invasion (4–6). Although there has been a steady decrease in the malaria burden, particularly for *P. falciparum* (Pf) in the last decade, the impact is much less pronounced for Pv because of latent infections and greater transmissibility in Pv endemic areas (4). In addition, this trend has stagnated recently because of political and economic instability and the global health crisis caused by the COVID-19 pandemic (7–9). To address the burden of Pv malaria, additional strategies are needed.

Pv initiates blood-stage infections by invading immature red blood cells (RBCs), or reticulocytes using its endogenous Duffy binding protein (PvDBP) to access the Duffy antigen receptor for chemokines (DARC) (encoded by gene atypical chemokine receptor 1, ACKR1 (10)) (11–13). The structural biology for the PvDBP and DARC interaction has become increasingly well defined (14–18). Among six distinct structural regions, the cysteine-rich region II (PvDBPII) contains three subdomains (SD) (12). SD2 contains the Duffy antigen binding motif (10). As PvDBPII is the most polymorphic region, it is suggested to be under strong selection pressure (19, 20). This motif interacts with the N-terminal 30 amino acid region of DARC to form the heterotetramer necessary for the binding interaction and commitment to subsequent invasion (15). The necessity of PvDBPII for Pv invasion of reticulocytes makes it a primary target for host immunity. Evidence supporting this hypothesis has included observations of extensive amino acid variation (most highly abundant in SD2 and SD3) (21–23) and recent Phase I/IIa vaccine efficacy against blood-stage Pv infection (24).

We and others have produced murine monoclonal antibodies to PvDBPII that blocked the binding of DBPII to DARC in various binding inhibitory assays (25, 26). However, they were not strain-transcending and failed to inhibit Pv invasion of reticulocytes *in vitro* (25–27). These murine mAbs recognized SD3 of PvDBPII, which does not contain the binding motif to DARC, but it is important for developing the heterotetramer necessary for stable binding interaction to DARC (14, 15). In comparison, we have previously identified 9 to 15% of individuals living in Pv endemic areas of Papua New Guinea, Cambodia, and Brazil with antibodies to PvDBPII capable of blocking PvDBPII from binding to DARC and preventing Pv invasion of reticulocytes (28–33). High levels of these blocking antibodies correlate with reduced risk of infection and disease in human cohort studies (28–33). From some individuals with binding inhibitory antibodies to PvDBPII, we have isolated PvDBPII-specific memory B cells from which we have generated a panel of human monoclonal antibodies (humAbs) (33). Three of these humAbs have been tested in a short-term ex vivo growth assay using clinical Pv isolates from Cambodia and Brazil. Notably, these humAbs exhibited strain-transcending inhibition of Pv reticulocyte invasion by up to 80% at 100 μg/mL and two of these humAbs recognized the predicted DARC binding site in PvDBPII SD2 (22, 27, 33).

Additional humAbs have been generated from healthy volunteers immunized with a vaccinia virus vectored vaccine expressing Salvador I strain (Sal I) PvDBPII in a Phase Ia clinical vaccine trial (34). The humAbs from this vaccine trial were validated in recombinant PvDBP-DARC binding inhibition assays, *ex-vivo* Pv invasion assays, and *P. knowlesi* (Pk) growth inhibition assays, using the Pk strain A1-H.1 PvDBP OR /Δ14 (PkPvDBPOR), which has been CRISPR-Cas9 modified to replace Pk’s endogenous DARC binding protein with PvDBP and adapted to grow in continuous human culture (35–38). Several of these humAbs displayed strain-transcendent blocking of recombinant PvDBPII to the DARC ectodomain and inhibited invasion, including an SD3-specific humAb (38). Here, we built upon this research by using the PkPvDBPOR *in vitro* model system to examine inhibition of human erythrocyte invasion by naturally acquired PvDBPII-specific humAbs (33). We expand the analysis of vaccine-induced and naturally acquired humAbs, individually and in combination, to identify potential additive and/or synergistic effects.

## RESULTS

### Characteristics of individual humAb in the PkPvDBPOR Growth Inhibtion Assay (GIA)

To examine and compare the functional activity of the NA and VI humAbs, we developed a modified GIA with PkPvDBPOR (Fig.1). The assay requires the addition of CellTrace-labeled, enriched parasitized cells (donor) to uninfected cells (recipient) to ensure identification of newly established invasion of recipient target cells as opposed to identification of infected donor cells from the feeder culture. This experimental design optimized the detection of new erythrocyte invasion events and was the basis of our assessment of humAb inhibition characteristics.

**FIG 1.**
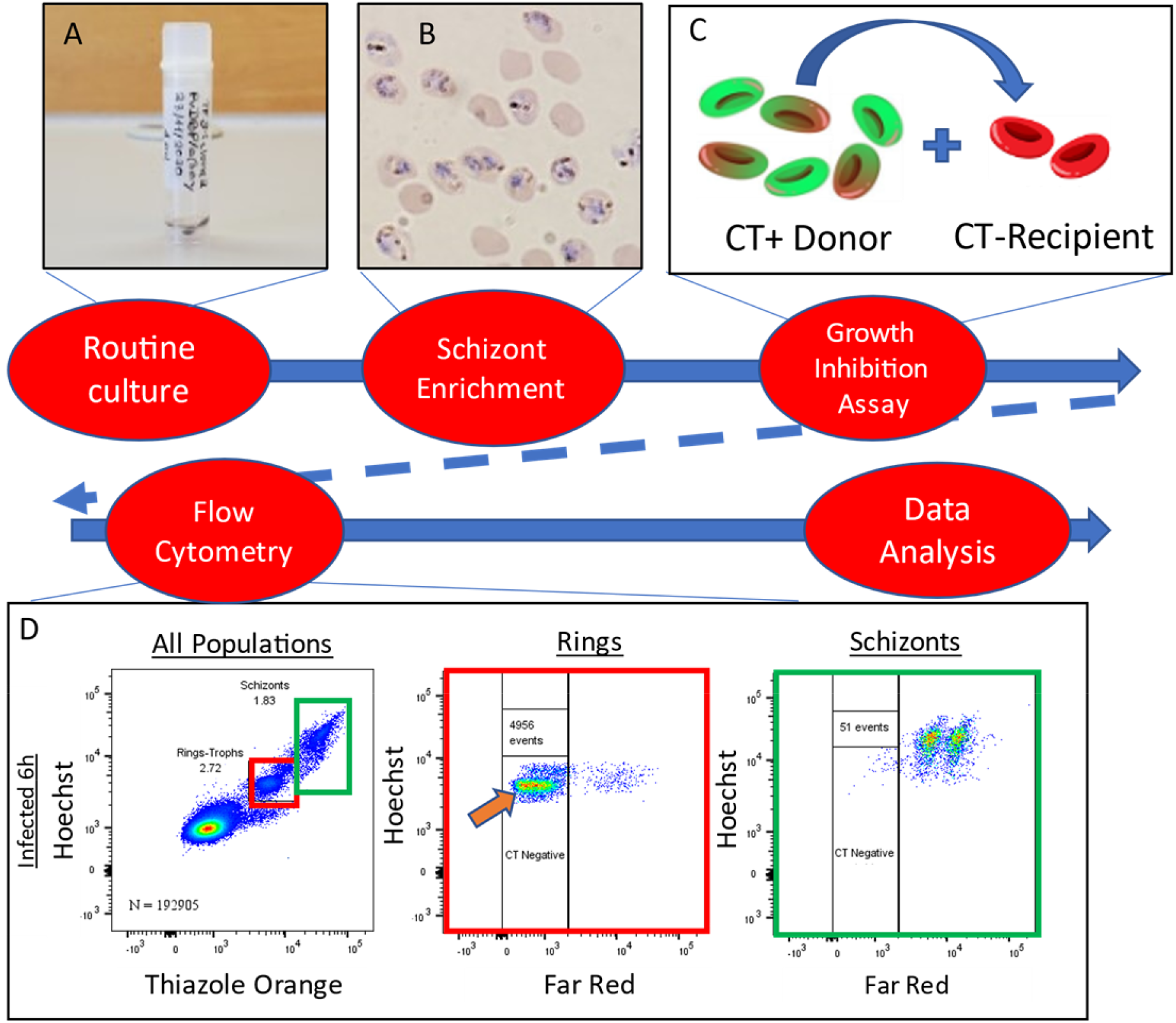
Experimental Design: **(A)** Cultures were initiated from a cryopreserved stock of *P. knowlesi* A1.H.1 strain and were maintained in human RBCs at 2% hematocrit. **(B)** Parasites were enriched for schizonts using a Nycodenz gradient. (C) The schizont preparation was labeled with CellTrace to identify them as donor cells. Donor cells were then mixed with unlabeled recipient cells at a ratio of 1:20 and incubated with/without experimental reagents. (D) After 6 hours of culture, samples were stained with Hoechst 33342 for DNA content (Y-axis, and thiazole orange, left-hand panel and Far Red (CellTrace, middle and right panels) to identify rings (red boxes) that represent new invasion events (CT negative, N=4956) and schizonts (green boxes).

After mixing the labeled donor cells with unlabeled recipient cells, measurement of samples at baseline (time zero) revealed up to 0.16% ring stage parasites and 3.03% schizonts (Fig. 2A, upper panels). Greater than 99.9% of schizonts were labeled with CellTrace whereas 88 events of ring-stage parasites were CellTrace negative (Fig. 2A, upper middle and right columns). The CellTrace negative cells with ring-stage parasites likely represent early infection of newly added recipient target cells. The number of newly infected cells that are CellTrace negative markedly increased 55-fold (4956 events) by 6 hours (Fig. 2A, second row) in the absence of antibodies or with a non-PvDBP-specific control humAb 043048 (tetanus toxin C-terminal fragment-specific humAb; Fig. 2A, lower row). The ring-stage parasite observed at time zero are subtracted from new invasion events in the 6-hour cultures in different experimental conditions. To demonstrate that PkPvDBPOR invasion of human red cells is Duffy dependent, the nanobody CA111 (that recognizes an epitope on DARC to which PvDBPII binds) inhibited PkPvDBPOR invasion by 91% (Fig. 2A). The NA PvDBPII humAb 099100 is a focal point for our comparisons in the PkPvDBPOR model as it was found to have the highest avidity in our earlier studies and shown to consistently inhibit Pv clinical isolates in vitro (33, 39). As shown in Fig. 2B and C, 099100 inhibited PkPvDBPOR erythrocyte invasion in a dose-dependent fashion up to 85% and showed an IC_50_ of 135 μg/mL. In addition to 099100, eight additional NA humAbs were tested and analyzed for growth inhibition of PkPvDBPOR (Fig. 3A). Their IC_50_ values were aggregated into a summary curve, ranging from 50.7 μg/mL to 338.4 μg/mL, all with R^2^ values above 0.95 (Fig. 3B and D). The NA humAb 065098 best inhibited PkPvDBPOR transgenic parasites with an IC_50_ of 50.7 μg/mL.

**FIG 2.**
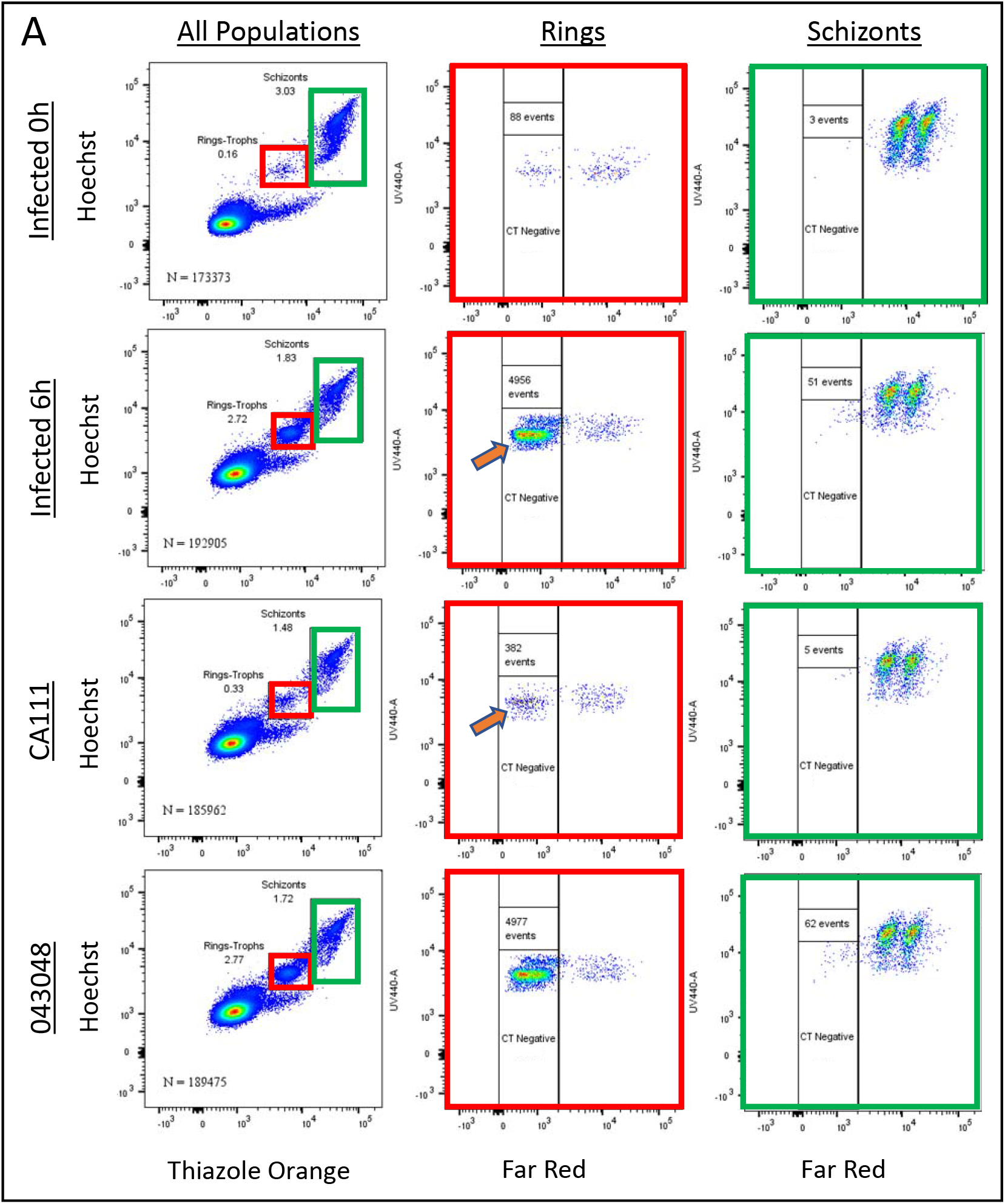
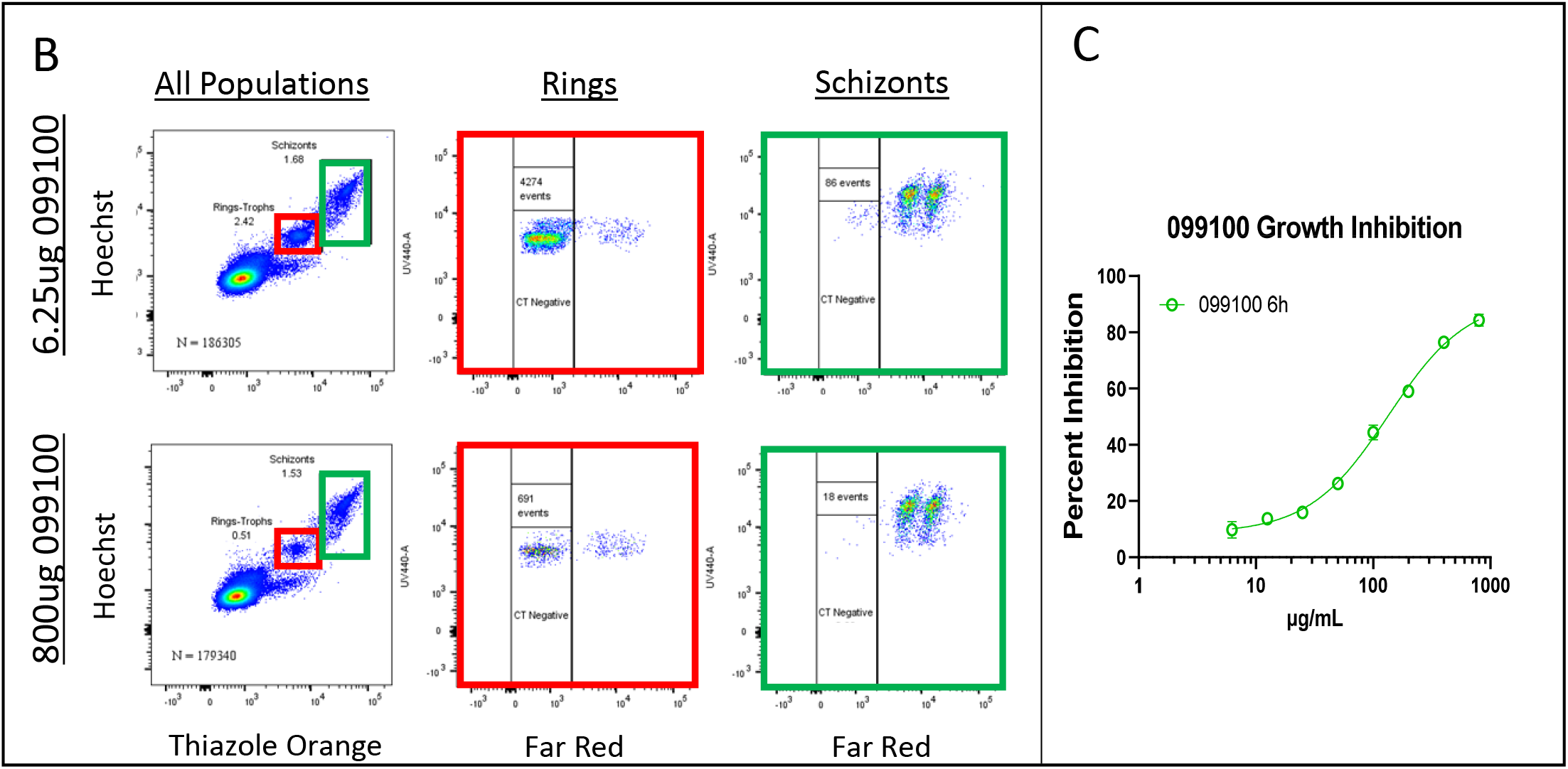
Invasion inhibition activity for humAb 099100 to PvDBPII: (A) Rows one and two show cultures at time zero hour and at 6 hours respectively, indicating new invasion events (orange arrow). Row three shows the blocking of PkPvDBPOR invasion with a camelid nanobody CA111. Row four shows culture containing a negative control humAb 043048 (tetanus toxoid-specific). **(B)** The top and bottom rows show 6 hour cultures treated with humAb 099100 at the lowest and highest concentration, respectively, of a two-fold dose response. **(C)** The full growth inhibition curve of 099100.

**FIG 3.**
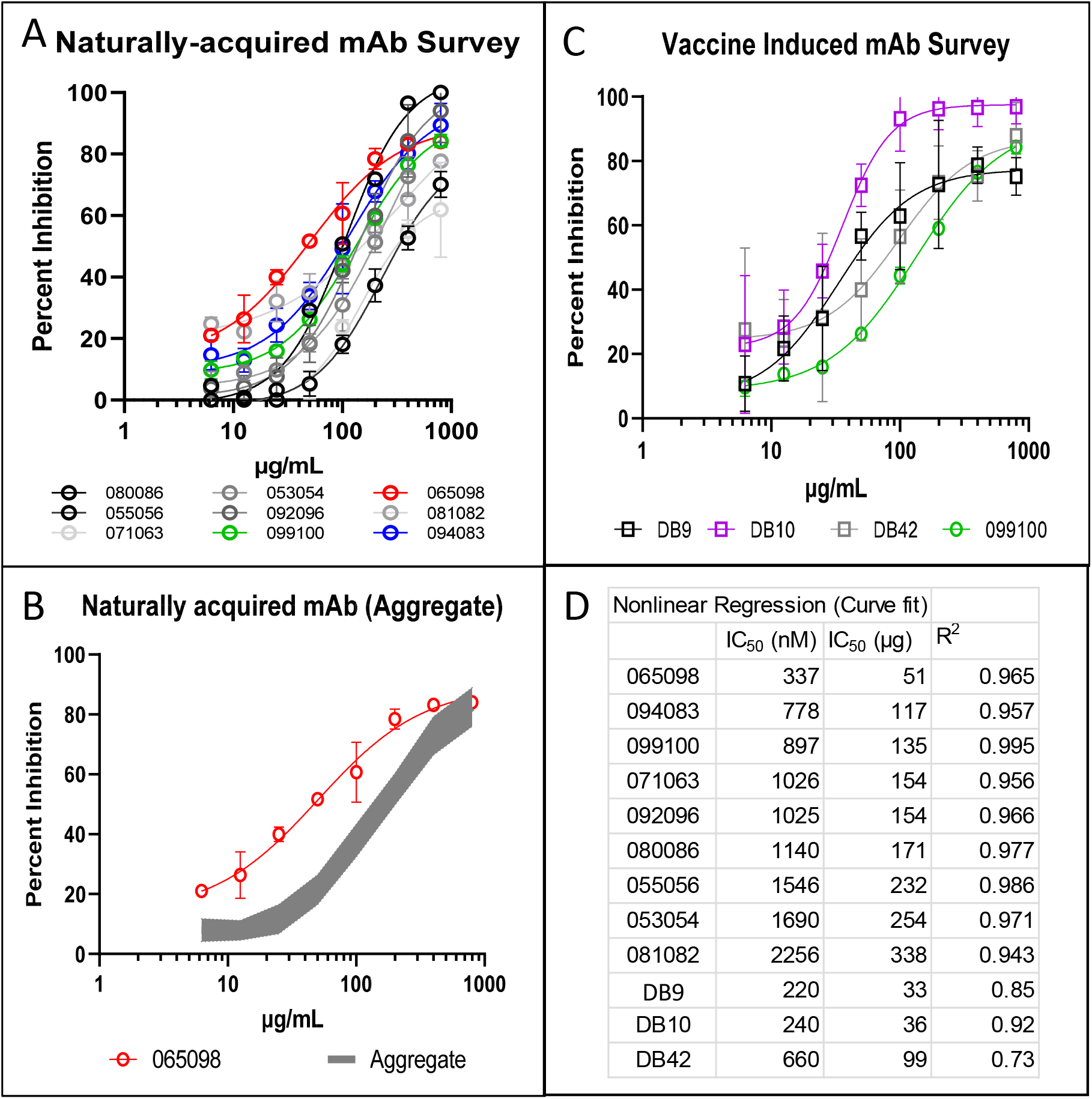
Invasion inhibition survey of all humAbs: **(A)** The full growth inhibition curves of 9 humAbs. **(B)** Eight humAbs aggregated into one curve with IC_50_ values ranging from 117 μg/mL to 338 μg/mL (gray curve); the best inhibitory humAb 065098 with an IC_50_ value of 51 μg/mL. **(C)** The full growth inhibition curves for 3 vaccine-induced humAbs; 099100 (green curve) overlaid for representative comparison to naturally-acquired humAbs, data referenced from panel A. **(D)** Nonlinear regression analysis to calculate IC_50_ values and goodness 2 of fit (R^2^ values) for all humAbs tested. All percent inhibition points represent mean and standard deviation of triplicate cultures.

We then evaluated how the VI humAbs, derived from humans vaccinated with the Sal 1 PvDBPII vaccine formulation (38) inhibited erythrocyte invasion and growth of PkPvDBPOR. Results for the independent tests of these three humAbs, DB9, DB10, and DB42 are summarized in Fig. 3C. These VI humAbs were characterized by IC50 values of 33.0 μg/mL, 35.6 μg/mL and 99.1 μg/mL (DB9, DB10, and DB42, respectively). Despite having lower IC_50_ values than 099100,DB9 (targets PvDBPII SD3 epitope (38)) and DB42 (PvDBPII subdomain epitope unknown) performed comparably to 099100, whereas the IC_50_ for DB10 was 4-fold lower than 099100 and nearly reached 100% invasion inhibition at 100 μg/mL (Fig. 3C).

### humAb combinations in the PkPvDBPOR GIA

Finally, we investigated whether combining two humAbs may have synergistic effects. Results presented here, focused on three specific humAbs, 099100, 094083, and 065098; based on distinct inhibition curves and different predicted PvDBPII binding epitopes from previously performed competition experiments (33). HumAb 065098 demonstrated the most potent GIA effect of the NA antibodies. The NA humAbs 092096, and 053054 bind to PvDBPII-DARC binding interface assessed by X-ray crystallographic studies (14, 15)) and exhibited competitive binding with 099100. This suggest that 099100 may bind to the same or nearby epitopes (33), although this does not exclude the possibility that they recognized different, but overlapping epitopes. Based on an absence of competitive binding with other humAbs, 094083 appeared to bind to a unique epitope (33). We also performed these combination studies with all three VI humAbs.

In these humAb combination studies, GIAs were performed with and without 099100 at its observed IC_25_ (50 μg/mL). Concentrations of the paired humAbs (065098 and 094083) were diluted two-fold starting at 800 μg/mL down to 6.25 μg/mL (Fig. 4A and C). Both 065098 and 094083 display increased levels of inhibition when combined with 099100, particularly 094083 (Fig. 4). The humAb 065098 showed marginal additivity (BSS of 1.64) with a peak at 100 μg/mL (Fig. 4B). The combination of 094083 and 099100 exhibited stronger additivity (BSS of 6.84; BSS of 9.9 judged to indicate synergy), and an expanded peak from approximately 12.5 μg/mL to 200 μg/mL, suggesting additivity at a broad range of concentrations.

**FIG 4.**
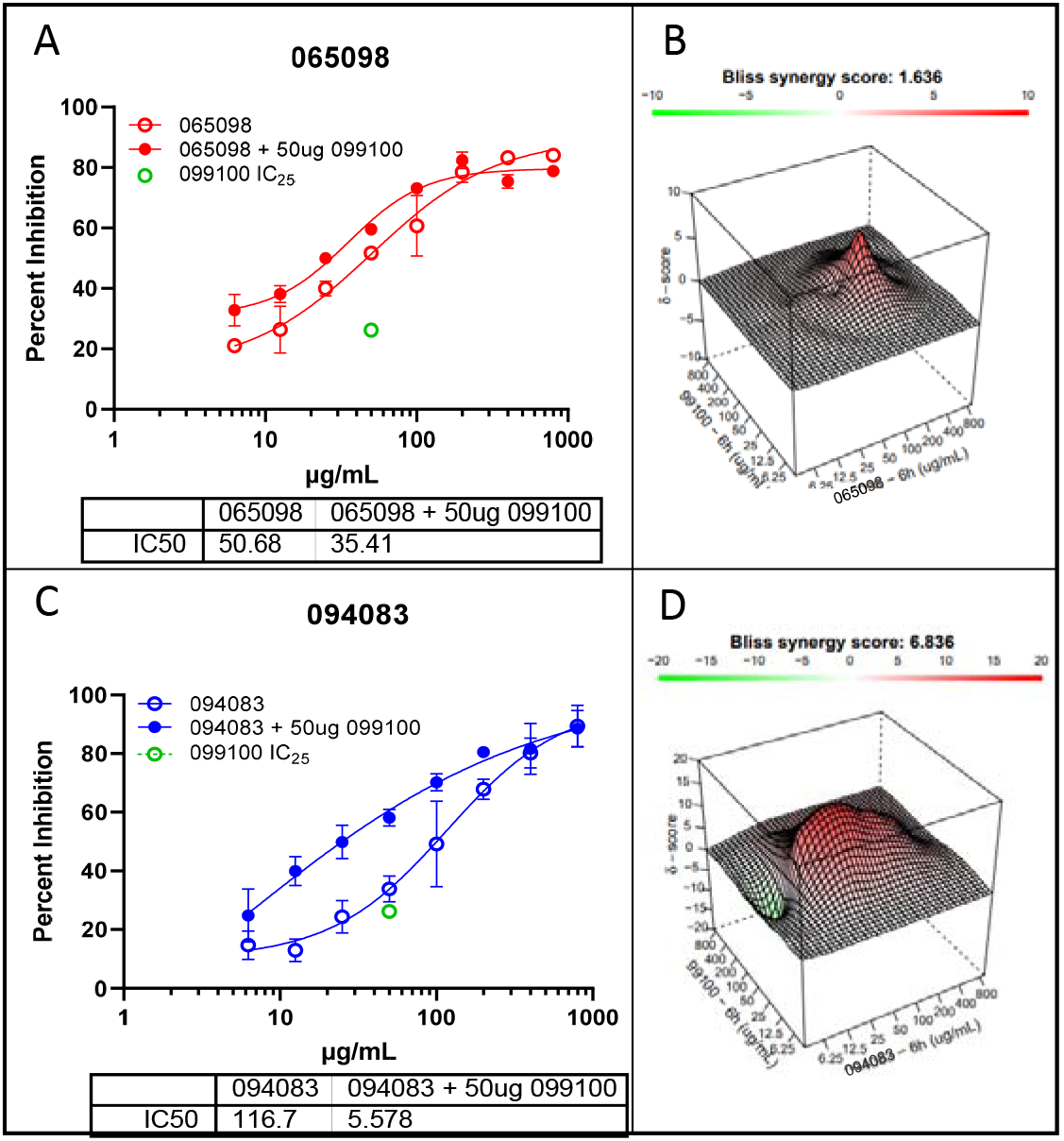
HumAb Synergy Experiments: **(A, C)** The first column shows growth inhibition curves for humAbs 065098, and 094083 individually, with and without the presence of 099100; 099100 was held constant at its IC_25_ (50 μg/mL) at each concentration (i.e., 800 μg/mL 065098 + 50 μg/mL 099100, 400 μg/mL 065098 + 50 μg/mL 099100, etc.); the green dot depicts the percent inhibition of 099100 at its IC_25_ (50 μg/mL); All points are mean and standard deviation of triplicate culture. The table below the figures shows differences in IC50 of 065098 and 094083 with and without 099100. **(B, D)** Assessment by SynergyFinder webbased application software to visualize multi-antibody multi dose combination data based on the independent model Bliss: 3-dimensional representation of the dose-response matrices showing varying concentrations of 065098 **(B)** and 094083 **(D)** on the x-axis, varying concentrations of 099100 on the y-axis, and synergy score (δ) on the z-axis; each matrix is color coded to show synergy distribution, a corresponding Bliss synergy score (BSS), and topography (peaks/valleys) at specific concentrations; green/valleys denotes antagonism, red/peaks denotes additivity or synergy. Scale for BSS: score<0 = antagonism, 10<score>0 = additivity, score>9.9 = synergy.

To evaluate whether humAbs generated by exposure to the PvDBPII vaccination might exhibit synergistic effects, we performed individual assessments of DB9 (of note DB is predicted to bind PvDBPII SD3), DB10 and DB42 (Fig. 5A, C and E), in combination with 099100. Relative to their individual inhibition curves, these humAbs produced antagonistic effects (Fig. 5B, D and F) when the concentration of monoclonals exceeded 25 μg/mL. The BSS for each combination was −12.03 (DB9+099100), −8.18 (DB10+099100), and −13.5 (DB42+099100). At lower concentrations of the VI humAbs, they showed additive effects; IC_50_ values for DB9 and DB10 in combination with a fixed amount of 099100 had increased to 299.4 μg/mL and 73.9 μg/mL respectively (Fig. 5A and C). The values generated in the DB42+099100 combination did not generate a distinct inhibition curve, therefore a stable IC_50_ could not be calculated for DB42 (Fig. 5E).

**FIG 5.**
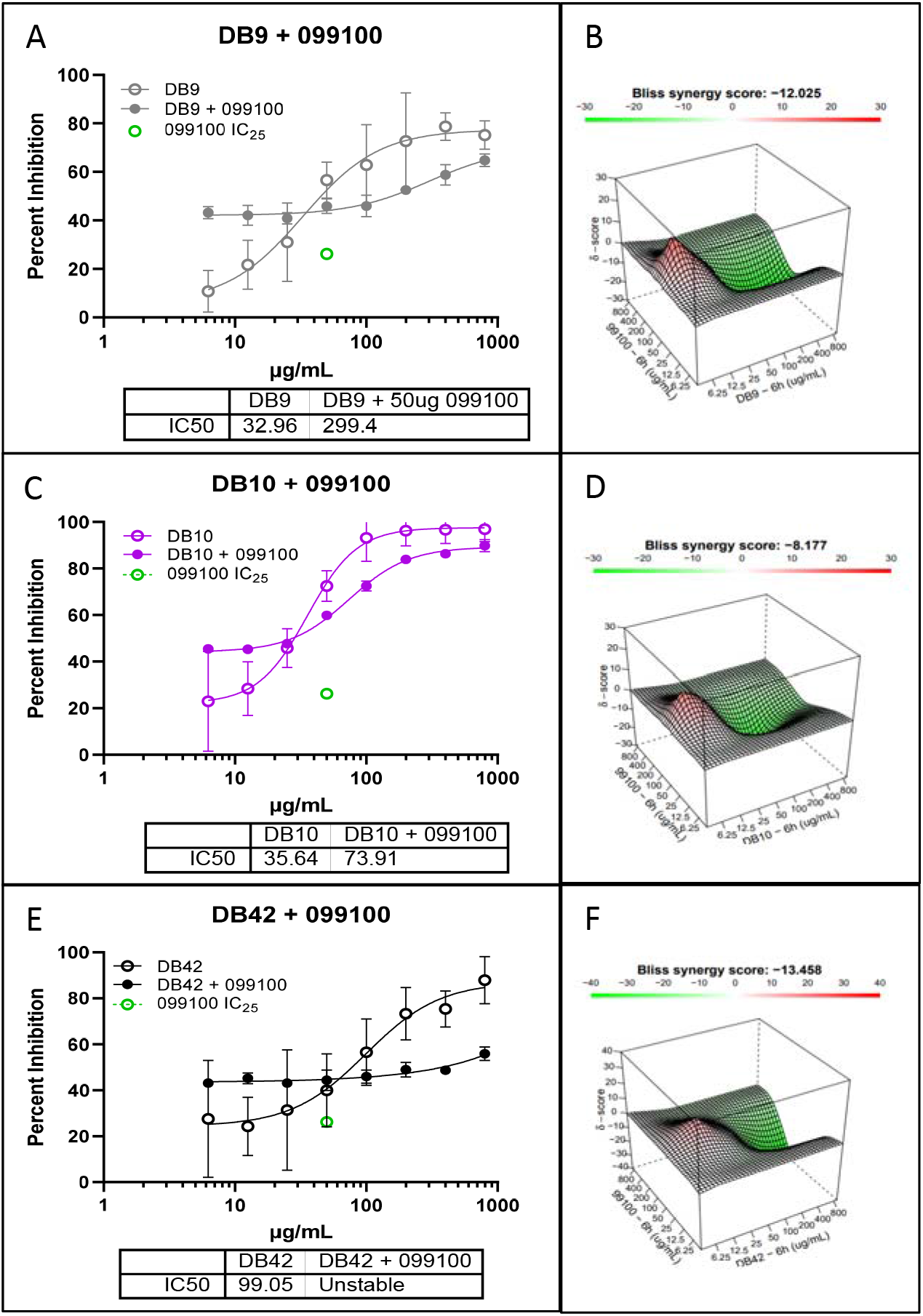
Invasion inhibition with combination of humAb 099100 with humAbs DB9, DB10 and DB42: **(A, C, E)** Growth inhibition curves for humAbs DB9, DB10, DB42, and 099100 individually, with and without the presence of 099100 held constant at its IC_25_ (50 μg/mL)(green dot). All points represent mean and standard deviation with triplicate cultures. **(B, D, F)** 3-dimensional representations of the dose-response matrices showing varying concentrations of DB9 **(B)**, DB10 **(D)**, and DB42 **(F)** on the x-axis, respectively from top to bottom, 099100 on the y-axis, and synergy score (δ) on the z-axis. Refer to legend in Figure 4 describing Bliss synergy matrix and scores.

The best blocking humAbs that target epitopes within PvDBPII SD2, NA 065098 and VI DB10, were examined in combination as to whether they would generate a synergistic response (Fig. 6). The combination experiments were performed reciprocally, with one antibody being varied across a two-fold dose-response range (eight concentrations - 800 μg/mL to 6.25 μg/mL) with the other being fixed at 50 μg/mL. These combination experiments had no additive or synergistic effect (Fig. 6A), as inhibition values were only marginally different from the independent inhibition dose responses. IC_50_ values for DB10+065098(fixed concentration) increased from 35.6 μg/mL to 64.99 μg/mL and decreased from 50.68 μg/mL to 48.33 μg/mL for the 065098+DB10(fixed concentration). According to the synergy distribution and tensor, there was a strong antagonism between the two antibodies, resulting in a BSS of −25.2 (Fig. 6B).

**FIG 6.**
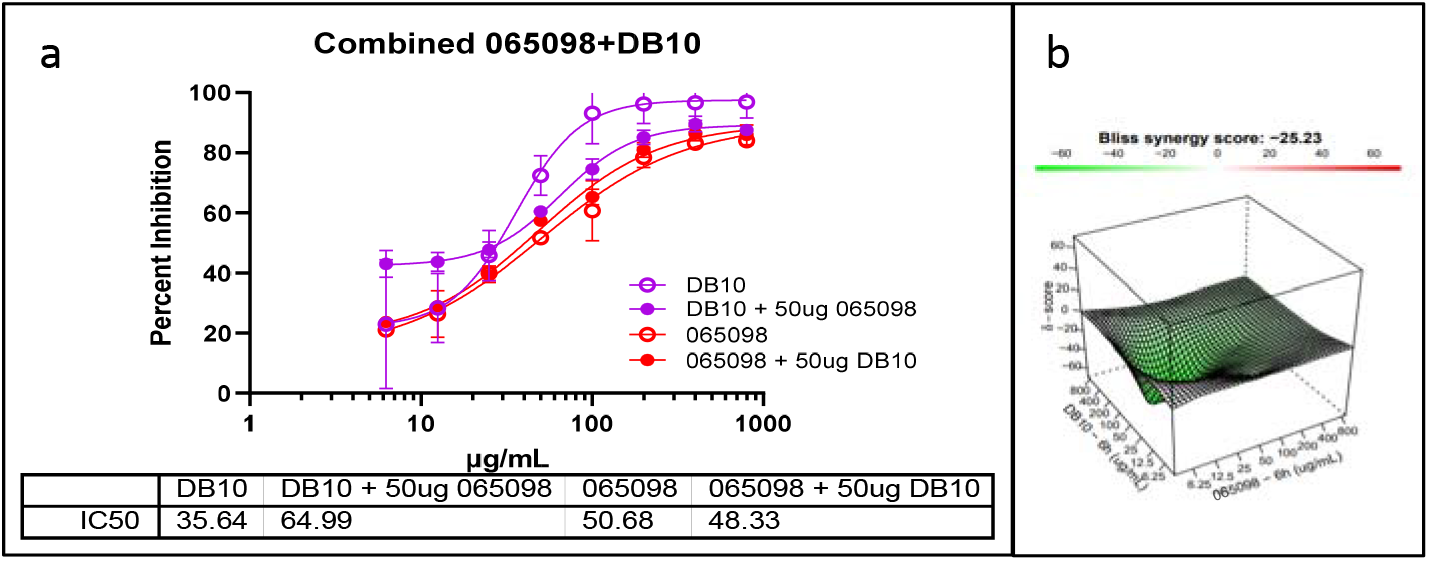
Combination experiment with humAbs 065098 and DB10: **(A)** Growth inhibition curves for humAbs DB10 and 065098 individually and in combination. One antibody was varied in a two-fold dose response at eight concentrations while the other was held constant at 50 μg/mL, and then inversely (red and violet curves); individual curves for DB10 (dashed blue curve) and 065098 (dashed green curve) are overlaid for comparison, data referenced from Fig. 3 and Fig. 4. All points represent mean and standard deviation with triplicate cultures. **(B)**3-D dose-response matrices showing the concentrations of 065098 on the x-axis, DB10 on the y-axis, and synergy score (δ) on the z-axis. Refer to legend in Figure 4 describing Bliss synergy matrix and scores.

## DISCUSSION

This study used PvDBPII-specific humAbs from naturally infected and vaccine-exposed people (21, 22) to inhibit human erythrocyte invasion by Pk genetically modified to express the Sal I allele of PvDBP, PkPvDBPOR (32–34). The results of our *in vitro* studies demonstrate an efficient approach for evaluating these antibodies individually and in combination. Results suggest that this *in vitro* system may help optimize the development of humAb prophylaxis and treatment strategies against blood-stage *P. vivax* malaria.

Our recent studies characterized these PvDBP-specific humAbs in the context of binding inhibition analyses (33, 38), *in vitro* inhibition (*ex vivo* Pv and *in vitro* Pk) (33, 38) and X-ray crystallography illustrating protein-protein binding specificities (27, 38). Here we have used the PkPvDBPOR *in vitro* system to perform individual invasion inhibition studies in side-by-side comparisons of the NA and VI PvDBP-specific humAbs. Additionally, based upon a rationale using humAbs with known affinity and binding site characteristics, we have tested hypotheses regarding specific interactions between these PvDBPII-specific humAbs (and the target antigen) to test the potential for additive, synergistic or antagonistic effects.

In tests on humAbs individually, we observed that the IC_50_ values for NA humAbs ranged from 51 to 338 μg/mL (340 - 2250 nM). In comparison, the vaccine-induced humAbs were three of the four best inhibitors with IC_50_ values of 32.96, 35.64, and 99.05 μg/mL (219.73, 237.60, and 660.33 nM) for DB9, DB10, and DB42 respectively (Mann Whitney P-value: 0.0182); the IC_50_ for the DARC-specific camelid nanobody, CA111, was 0.25 μg/mL (17 nM). The increased efficacy of the VI humAbs is not surprising considering they were generated from individuals immunized with Sal I PvDBP and were tested against PkPvDBPOR containing the Sal I variant (38). In contrast the NA humAbs were generated from Cambodian donors (33). In Cambodia, the Sal I variant of PvDBP is present but is not as common, potentially introducing more epitope variation by the naturally acquired humAbs (23). Overall, the humAbs reached a maximum inhibition above 80% at 800 μg (5.3 μM); 80% inhibition for CA111 was 2 μg (133 nM). We also found that the binding avidity data did not necessarily correspond with the erythrocyte invasion inhibition data for PkPvDBPOR (e.g., 065098 and 094083 exhibited among the weaker avidities but had the strongest erythrocyte invasion inhibitory effects, based on lowest IC_50_s blocking PkPvDBPOR *in vitro* invasion).

The humAb combinatorial tests, provided new insights beyond earlier studies that begin to identify specific humAb partners that may be able to optimize the therapeutic use of these reagents. For example, in our earlier studies, 099100 binding to PvDBP was competitively inhibited by 065098, but not by 094083 – suggesting that these two humAbs bind to different epitopes. Interestingly, the combinatorial tests showed negligible additive inhibition by 099100 + 065098, but significantly more inhibition by the 099100 + 094083 combination. Following this logic, we hypothesized that combinations of PvDBPII SD2-binding humAbs with PvDBPII SD3-binding humAbs would demonstrate even greater additive, possibly synergistic inhibition. Instead, we observed that this combination (DB9+099100(fixed concentration)) rendered antagonistic effects on PkPvDBPOR invasion of human erythrocytes.

## CONCLUSION

Using monoclonal antibodies against viral and parasite invasion ligands has demonstrated how challenging the discovery of the optimal combination of reagents to block infection can be (40–46). The technologies applied in this and recent studies (33, 38) appear to expand the generation and evaluation of humAb therapeutic reagents for treatment and prophylaxis of *vivax* malaria that have not been possible previously.

These efforts continued the exploration of *in vitro* methods for the culture of *P. vivax.*The treatment of *vivax* malaria in permissive non-human primate models is an important next step for evaluating these potentially protective humAbs. For example, the humAbs may have greater in vivo activity enhanced by Fc-mediated activity. The further development of these methods and reagents will be important if *P. vivax* is to be eliminated as a significant global public health challenge.

## MATERIALS AND METHODS

### Ethics approval and consent to participate

All studies were conducted following protocols approved by the University Hospitals of Cleveland Institutional Review Board (#08-03-33 and #09-90-195).

### Human Blood Preparation

We collected venous blood from healthy consented donors in EDTA vacutainers. The blood was centrifuged (1500xg for 5 minutes) to separate plasma and cellular material. Plasma is aspirated, and the remaining blood is passed through a neonatal blood filter (Haemonetics NEO1) for leukocyte depletion, washed with PBS, and centrifuged (2000xg for 8 min). We aspirated PBS off. An equal volume of Pk complete medium (see below) is added to bring the blood to 50% hematocrit and stored at 4°C. Stored blood will support culture growth for approximately 3 weeks. Fresh blood is acquired every 2 weeks or earlier. The Duffy (Fy) genotype was assessed as previously described (47). Donors for Pk culture were either Fy A+/B+ or Fy B+/B+.

### Monoclonal antibodies

Cloning, expression and purification of nine human PvDBP-specific monoclonal antibodies (humAbs: 099100; 080086; 055056; 071063; 053054; 092096; 065098; 081082; 094083) has been previously described (33, 48, 49). Three PvDBP-specific humAbs (DB9; DB10; DB42) were generated from individuals exposed to a vaccinia virus vectored vaccine expressing Salvador I strain (Sal I) PvDBPII in a Phase Ia clinical vaccine trial (34). A humAb specific for tetanus toxoid C-terminal (043048) was used as a negative control. For these experiments protein concentration was determined using a Nanodrop at 280 nm. The mAbs were concentrated to >4 mg/ml and filter-sterilized through a 0.22 μm PVDF filter for subsequent use. As a control to demonstrate invasion inhibition of PkPvDBPOR we used a nanobody (CA111) specific to the Fy6 epitope on DARC that blocks the binding of PvDBP to DARC as previously described (50).

### *Plasmodium knowlesi in vitro* culture

Pk culture media (PkCM) included RPMI 1640 medium (22400, Gibco) supplemented with 1.15 g/L sodium bicarbonate, 1 g/L dextrose, 0.05 g/L hypoxanthine, 5 g/L Albumax II, 0.025 g/L gentamicin sulfate, 0.292 g/L L-glutamine, and 10% (vol/vol) heat-inactivated horse serum (26050, Gibco) referred to as PkCM as previously described (37). Pk cultures were maintained in sealed flasks with 5% O_2_, 5-7% CO_2_, balanced by nitrogen.

Cryopreserved isolates of Pk A1.H.1 strain (1 mL) were thawed by drop-wise addition of 3.5% (weight/vol) NaCl over 1 minute, then transferred to a 15ml Falcon tube. Thawed parasites were centrifuged (1500xg, 5 min), and the supernatant was discarded. We repeated this treatment with 3.5% NaCl for three times. The treated parasites were pelleted (1500xg for 5 min) and resuspended in 1 mL of warm PkCM. This resuspension was added to 50 mL PkCM plus 1 mL of fresh RBCs (2% Haematocrit) (Fig. 1A).

The parasitemia of routine cultures was maintained below 5% and expanded to between 8-12% for growth inhibition assays. Routine culture maintenance included medium changes/dilution of parasites every 2-3 days. Giemsa-stained culture smears were made during culture changes to monitor parasite viability. Pk cultures were expanded for a minimum of 4 life cycles, synchronized using Nycodenz gradient to enrich schizonts for growth inhibition assays.

### Nycodenz Synchronization

Parasites were synchronized using Nycodenz (157750, MP Biomedicals) (Fig. 1b) as described previously (37). Nycodenz is a non-ionic tri-iodinated derivative of benzoic acid (51). Nycodenz stock solution is prepared at 27.6% (weight/vol) Nycodenz, 10% (vol/vol) 100 mM HEPES (BP299, Fisher BioReagents), adjusted to pH 7.0, supplemented with sterile distilled H_2_O to reach final concentrations, and then filter sterilized. A Nycodenz working solution (55% vol/vol of stock solution) comprises 55 mL of Nycodenz stock solution to 45 mL of PkCM (without serum). Parasite cultures were centrifuged (1500xg, 8 min). The supernatant is aspirated, and the pellet is resuspended in 1 mL PkCM, to approximately 50% haematocrit, for a total volume of 2 mL. Parasites were layered over 5 mL Nycodenz working solution in a 15 mL conical tube and centrifuged (900xg, 12 min) with low brake/acceleration. The brown interphase containing schizonts is pipetted off and pelletized in a microcentrifuge (6000xg, 1 min). Pellet is washed once in 1 mL of PkCM, and once in 1x PBS before CellTrace staining. Immediately following Nycodenz enrichment, direct blood smears demonstrated 50-60% schizonts, and 1-2% ring stage or trophozoites.

### Growth Inhibition Assays (GIAs)

The Nycodenz-enriched schizonts (donor cells) were stained in CellTrace Far-red (C34564, Invitrogen) at 4.5μM in 1x PBS for 30 minutes at 37°C on a rotator in darkness. Donor cells were microcentrifuged (4000xg, 1 min), the supernatant aspirated, and the pellet was resuspended in 5 mL of PkCM for the growth inhibition assay. PkCM (50 μl) with humAbs were first aliquoted in 96 well flat bottomed microwell plates. To reach a final culture volume of 100 μl cultures, 25 μl of 8% RBCs (unlabeled recipients) in PkCM (final hematocrit of 2%) was added, followed by 25 μl of donor cells for a 1:20 ratio (donor:recipient). An additional well was made for the 0h time point, which was taken and fixed for starting parasitemia and time zero invasion events. The remaining enrichment preparation was fixed to verify CellTrace labeling efficiency by flow cytometry. The 96 well plates were placed in a modular incubator chamber (MIC-101), gas for 2 minutes (5% O_2_, 7% CO_2_, balanced by nitrogen), sealed and incubated in a 37°C incubator for 6 hours. At the end of the culture period, experimental samples were fixed in a 1x PBS solution containing 4% paraformaldehyde and 0.01% glutaraldehyde for 20 minutes at room temperature. Cells were centrifuged (4000xg, 5 min) and washed once with 1x PBS. Cells were stored in 1x PBS at 4°C or immediately stained for subsequent flow cytometry (Fig. 1c).

### Flow Cytometric Evaluation of GIAs

Samples were stained in a 1x PBS solution containing Hoechst 33342 at 4μM for DNA content (parasites) and thiazole orange at 100 ng/mL, for preferential staining of RNA and reticulate matter in reticulocytes, for a minimum of 30-40 minutes at room temperature, or overnight at 4°C. Samples are then monitored and analyzed by flow cytometry using Biosciences BD LSR II Flow Cytometer. The Hoechst dye was excited by the ultraviolet 355 nm laser (excitation peak 355 nm, emission peak 465 nm) and detected using a 440/40 filter. Thiazole orange was excited by a blue 488 nm laser (excitation peak 514 nm, emission peak 533 nm) and detected using a 525/20 filter. CellTrace Far-red was excited by the red 640 nm laser (excitation peak 630 nm, emission peak 661 nm) and detected using the R660/20 filter. Resulting FCS files were analyzed by Flowjo 10.8.1 software for growth inhibition (Fig. 1d). The percent growth inhibition was calculated based on a comparison between the CellTrace negative Ring gate at 6 hours experimental invasion events divided by 6h control invasion events, both minus zero-hour events.

### Statistical Analysis

We applied a nonlinear regression curve analysis to estimate the IC_50_ (antibody potency) and R^2^ values using GraphPad Prism 9 software. The Synergy Finder 2.0 web-based application software evaluated the synergistic, additive or antagonistic effects of a combination of two humAbs based on the independent model Bliss (52). Synergy Finder generates a 3-dimensional representation of the dose-response matrices showing the concentrations of one humAb on the x-axis, a second humAb on the y-axis, and the synergy score (δ) on the z-axis; each matrix is color coded to show synergy distribution, a corresponding Bliss synergy score (BSS), and topography (peaks/valleys) at specific concentrations; green/valleys denotes antagonism, red/peaks denotes additivity or synergy. Scale for BSS: score<0 = antagonism, 10<score>0 = additivity, score>9.9 = synergy.

## ACKNOWLEDGEMENTS

We thank the study donors who participated over the course of this investigation. We thank Olivier S. Bertrand (INSERM/University Paris Diderot) for the gift of anti-Fy antibody (CA1110). We thank Robert W. Moon for providing the PkPvDBPOR strain used in this study. RWM was supported by the UK Medical Research Council (MRC Career Development Award to RWM MR/M021157/1). This study was funded by grants from the Veterans Affairs Research Service (BX001350) and NIH Grants (AI064687 and AI143694) to CLK and an NIH Grant (AI097366 and AI148469) to PAZ. QDW was supported by the Immunology Training Program for pre-doctoral students (5T32AI089474; PI Brian Cobb).

